# Identifying Alzheimer’s disease-associated genes using PhenoGeneRanker

**DOI:** 10.1101/2024.11.12.623269

**Authors:** Most Tahmina Rahman, Fahad Saeed, Serdar Bozdag, Alzheimer’s Disease Neuroimaging Initiative

**Affiliations:** Department of Computer Science and Engineering, University of North Texas, 1155 Union Circle #311366 Denton, Texas 76203, United States; Department of Mathematics, University of North Texas, 1155 Union Circle #311366 Denton, Texas 76203, United States; BioDiscovery Institute, University of North Texas, 1155 Union Circle #311366 Denton, Texas 76203, United States; Center for Computational Life Sciences, University of North Texas, 1155 Union Circle #311366 Denton, Texas 76203, United States; Knight Foundation School of Computing and Information Sciences, Florida International University, 11200 SW 8 Street, CASE 354 Miami, Florida 33199, United States

**Keywords:** Alzheimer’s Disease genes, network biology, PhenoGeneRanker, network propagation, gene prioritization, APP

## Abstract

Alzheimer’s disease (AD) is a neurogenerative disease that affects millions worldwide with no effective treatment. Several studies have been conducted to decipher to genomic underpinnings of AD. Due to its complex nature, many genes have been found to be associated with AD. Despite these findings, the pathophysiology of the disease is still elusive. To discover new putative AD-associated genes, in this study, we integrated multimodal gene and phenotype datasets of AD using network biology methods to prioritize potential AD-related genes. We constructed a multiplex heterogeneous network composed of patient and gene similarity networks utilizing phenotypic and omics datasets of AD patients from the Alzheimer’s Disease Neuroimaging Initiative (ADNI) database. We applied PhenoGeneRanker to traverse this network to discover potential AD-associated genes. To assess the impact of each network layer and seed gene, we also run PhenoGeneRanker on different variants of the network and seed genes. Our results showed that top-ranked genes captured several known AD-related genes and were enriched in Gene Ontology (GO) terms related to AD. We also observed that several top-ranked genes that are not in AD-associated gene list had literature supporting their potential relevance to AD.

## 1 Background

Alzheimer’s disease (AD) is a complex and progressive neurogenerative disease, and is one of the leading causes of death among older people. AD has become a growing public health crisis, affecting about 50 million people around the world, and six million people in the US, leading to huge burden to patients, caregivers and the healthcare sector. By the year 2040, the entire direct medical expenses for AD are anticipated to reach $259 billion^1–3^. One of the hallmarks of AD is the accumulation of tau and amyloid-β (Aβ) proteins in the brain, which lead to neuron death^4,5^. Growing studies suggest that some of the biological factors of AD could occur decades before the symptoms of the disease are apparent^6,7^. Screening methods such as amyloid and tau position emission tomography (PET) could be effective for early AD detection^8^. Several computational methods such as VGG-TSwinformer^9^, PPAD^10^, TA-RNN^11^, and PVTAD^12^ have been developed for early prediction of AD, too.

To characterize AD biology, several consortiums such as Alzheimer’s Disease Neuroimaging Initiative (ADNI), Alzheimer’s Disease Sequencing Project (ADSP), and National Alzheimer’s Coordinating Center (NACC) have facilitated generation of vast amount on multimodal biomedical datasets such as transcriptomics, genomics, epigenomics (i.e., multi-omics), neuroimaging, and clinical. Genome Wide Association Studies (GWAS) have been conducted^13– 15^ to report several AD risk genes such as *APOE, APP, ABCA7, MPO, PSEN2*, and *PSEN1*, annotated in databases such as MedGen^16^, GTR^17^, PheGenI^18^, and AlzGene^19^. Further research studied these genes in more detail. For instance, increased level of *MPO* expression is associated with increased the risk of AD^20^. The *ABCA7* levels have been observed to be increased in the AD brain causing amyloid plaque burden^21^. Mutations within amyloid precursor protein (*APP*) might cause 10% to 15% of early-onset familial AD^22^. *APOE* has been identified as a genetic risk factor for both familial late-onset and sporadic late-onset AD in several studies. Potential driver genes were reported from transcriptomic data by identifying differentially expressed genes (DEG) between AD and normal samples and utilizing machine learning- or deep learning-based methods^23,24^.

Utilizing the growing amount of multimodal datasets for AD, new AD-associated genes could be identified. Gene prioritization techniques can be applied to rank these genes based on their potential relevance to AD. Such findings would help improve disease knowledge, diagnosis, and leads to effective treatment options to prevent, postpone, and eventually cure AD. Network-based methods are effective to rank genes with respect to their potential relevance to a phenotype (e.g., disease)^25^. Several network-based gene prioritization tools have been developed to prioritize disease related genes^26–28^. Among them, the Random Walk with Restart (RWR) that considered both global network information and closeness to the central node was utilized to prioritize genes. One of the drawbacks of RWR is its biasness towards high degree nodes^39^. Recently, PhenoGeneRanker^29^ was introduced, a tool to perform RWR to rank genes associated with a phenotype (e.g., disease). PhenoGeneRanker analyzes multiplex (i.e., having multiple edge types) heterogeneous (i.e., having multiple node types) networks, which consist of gene and phenotype networks. PhenoGeneRanker also generates empirical p-values for the ranking to address the degree bias issue of RWR.

In this study, we ran PhenoGeneRanker on a multiplex heterogeneous network for AD to rank genes based on their potential association to AD. Utilizing datasets from ADNI and protein-protein interactions (PPI) from STRING, we built gene and patient similarity networks and applied PhenoGeneRanker (Figure 1). We found some well-known AD genes in our top rankings. By doing the GO enrichment analysis of our significant-top 200 genes, we observed that GO terms were highly associated with AD-related GO terms. We also conducted a literature search for the top genes that are not in AD databases, and found several genes reported as AD-related. We also checked the importance of utilizing different data modalities for the gene rankings.

**Figure 1.**
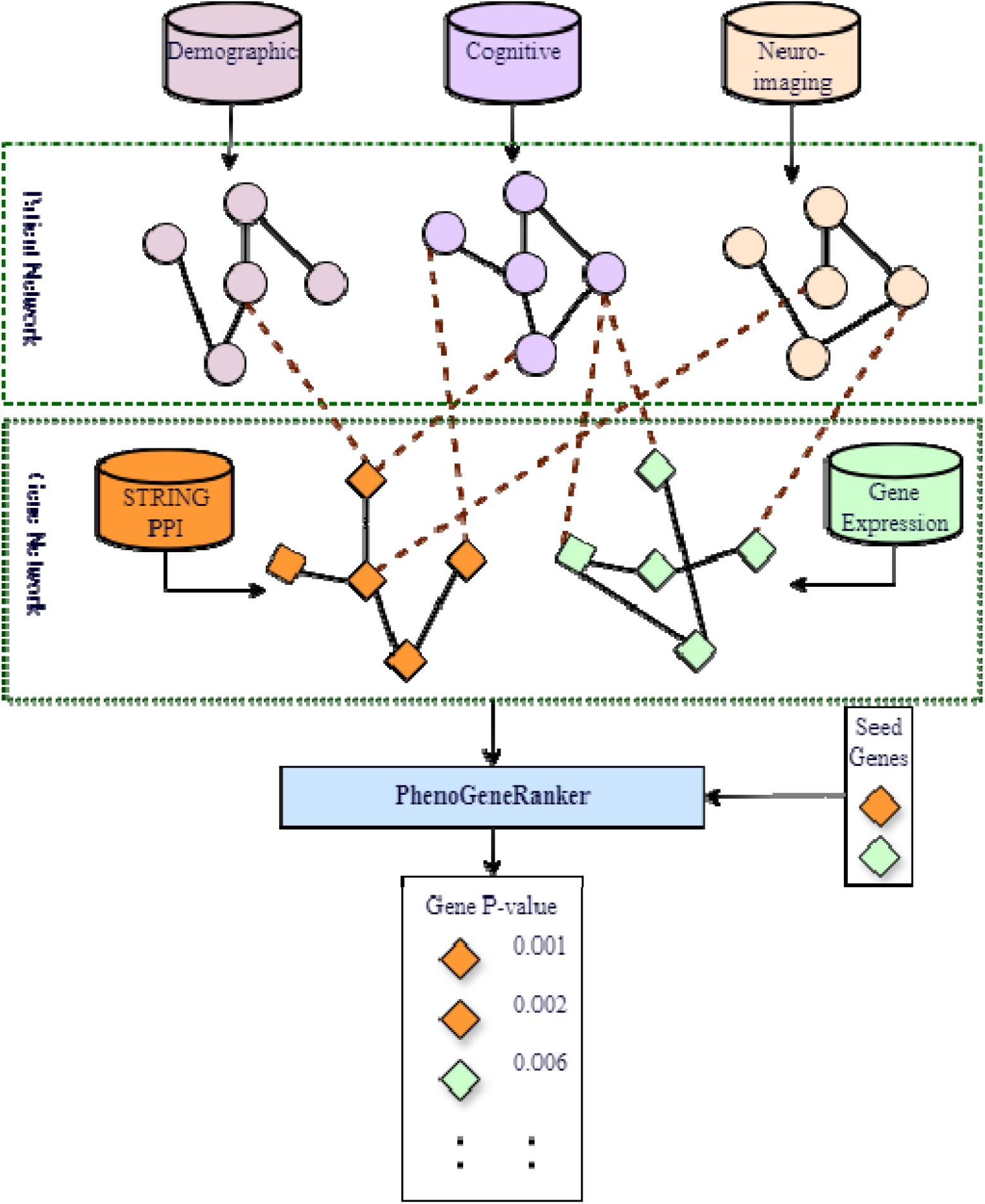
Overall pipeline to identify AD-associated genes. Three patient networks were generated from patients’ demographic, cognitive, and neuro-imaging data modalities. Two gene networks were built using gene expression and PPI data. PhenoGeneRanker was applied to the entire network to rank genes based on their potential association with AD seed genes. Finally, genes were ranked with associate p-value.

## 2 Methods

### 2.1 Building a multiplex heterogeneous network for AD

In this study, we built a multiplex heterogeneous network that consists of gene and phenotype node types and multiple relation (i.e., edge) types based on multiple data modalities. We obtained phenotypic and omics datasets of AD patients from the ADNI database^30^ (adni.loni.usc.edu). The ADNI was launched in 2003 as a public-private partnership, led by Principal Investigator Michael W. Weiner, MD. The primary goal of ADNI has been to test whether serial magnetic resonance imaging (MRI), positron emission tomography (PET), other biological markers, and clinical and neuropsychological assessment can be combined to measure the progression of MCI and early AD. Protein-protein interaction (PPI) dataset was collected from STRING database^31^ (https://string-db.org/). To make the network more AD specific, we relied on data from AD patients whenever available. Particularly, we obtained demographic, cognitive and neuroimaging data of AD patients using ADNIMERGE R package^30^ and gene expression data of AD patients from ADNI. In the ADNIMERGE dataset, most patients had longitudinal data. For these cases, we utilized the most recent data of each patient. For missing values, we used data from earlier time points to impute it.

We created three layers of phenotype network and two layers of the gene network. In the phenotype network where each node is an AD patient, the layers were built by computing similarity between AD patients based on demographic, cognitive, and neuroimaging data. The two gene layers were built based on gene expression and PPI datasets. We also built a bipartite layer to connect gene nodes to patient nodes based on expression activity of genes in AD patients. The features to build each layer are listed in Table 1.

**Table 1.**
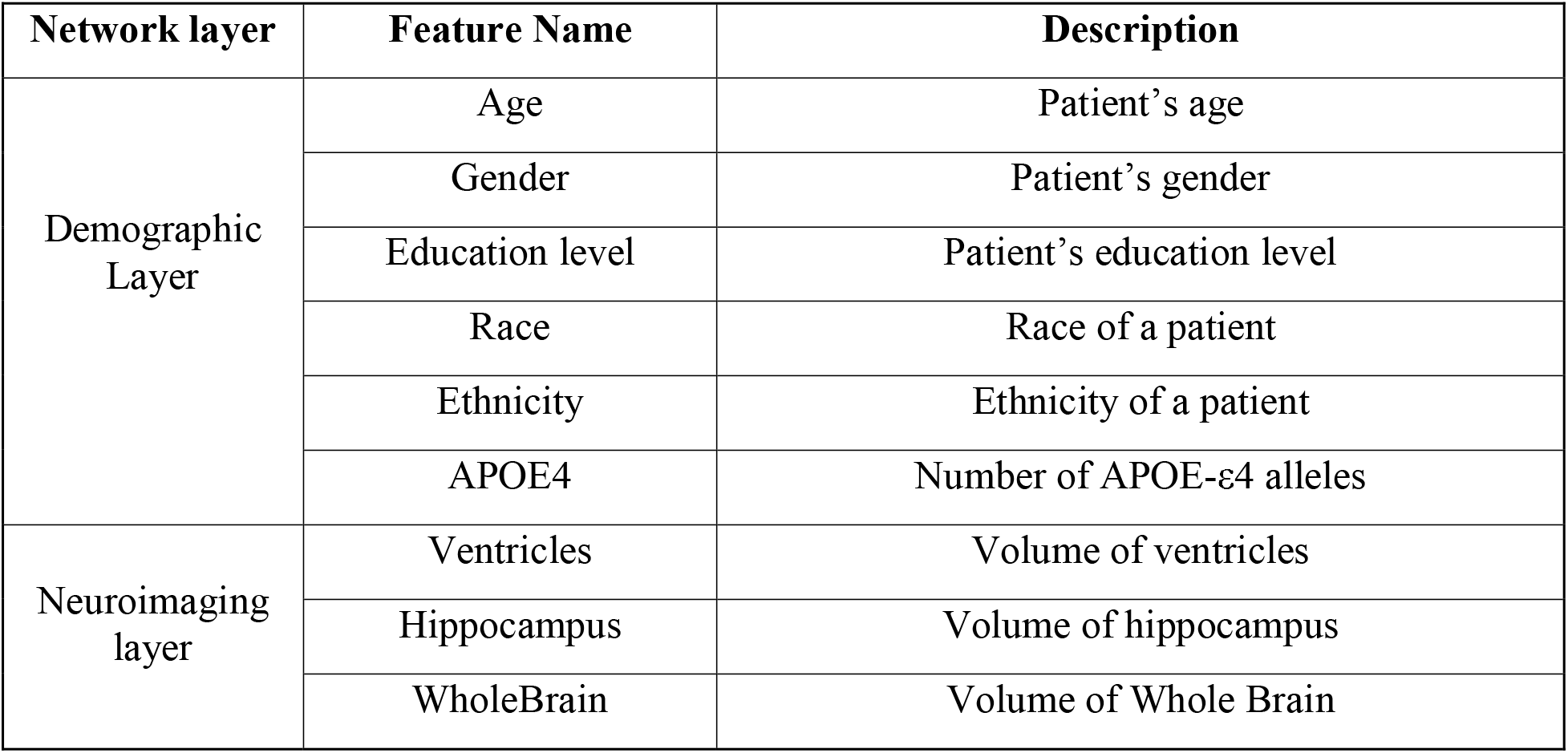
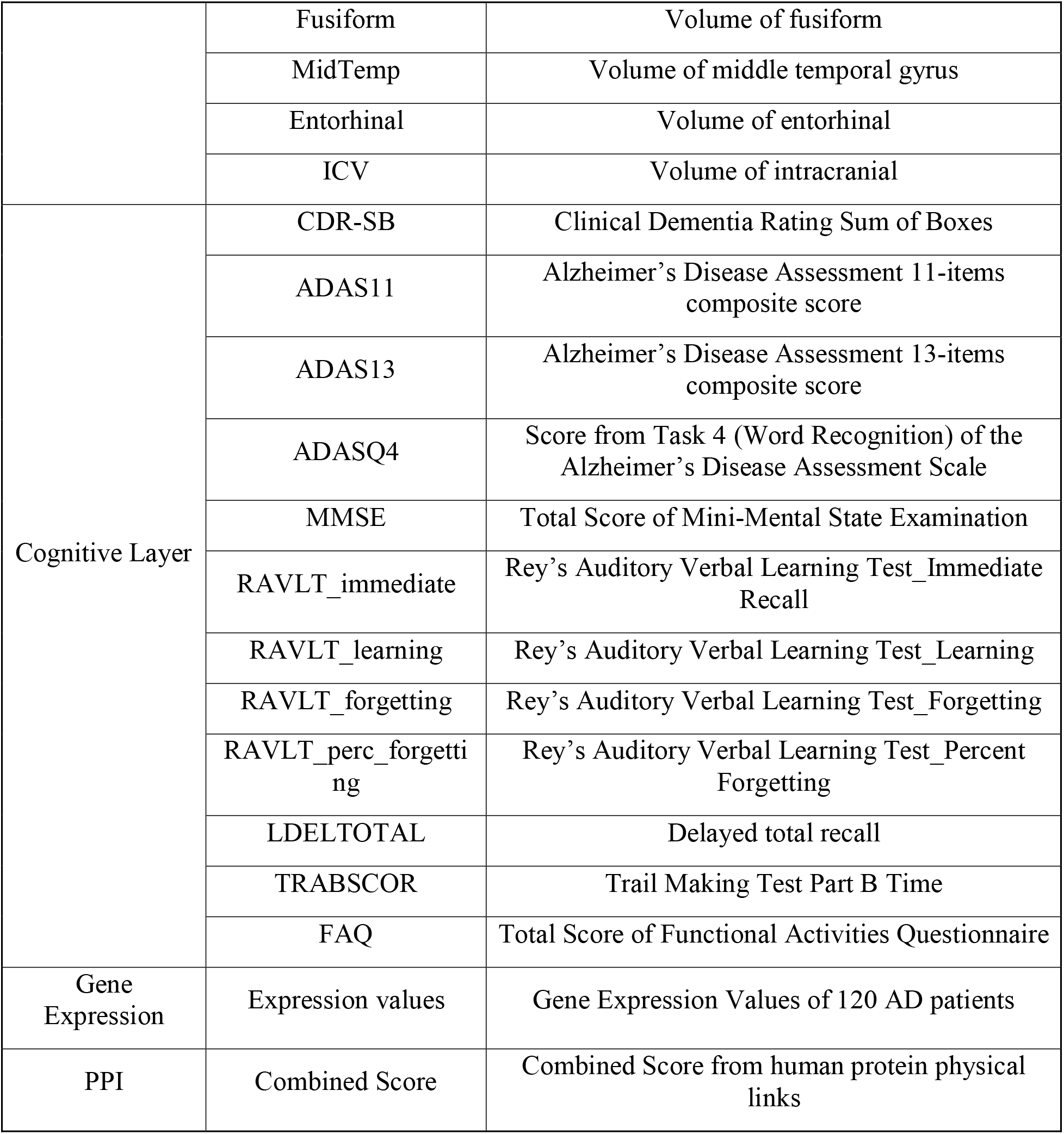
Details of features for each network from ADNI and STRING database.

| Network layer | Feature Name | Description |
| --- | --- | --- |
| Demographic Layer | Age | Patient's age |
|  | Gender | Patient's gender |
|  | Education level | Patient's education level |
|  | Race | Race of a patient |
|  | Ethnicity | Ethnicity of a patient |
| | APOE4 | Number of APOE-ε4 alleles |
| Neuroimaging layer | Ventricles | Volume of ventricles |
|  | Hippocampus | Volume of hippocampus |
|  | WholeBrain | Volume of Whole Brain |
|  | Fusiform | Volume of fusiform |
|  | MidTemp | Volume of middle temporal gyrus |
|  | Entorhinal | Volume of entorhinal |
|  | ICV | Volume of intracranial |
| Cognitive Layer | CDR-SB | Clinical Dementia Rating Sum of Boxes |
|  | ADAS11 | Alzheimer's Disease Assessment 11-items composite score |
|  | ADAS13 | Alzheimer's Disease Assessment 13-items composite score |
|  | ADASQ4 | Score from Task 4 (Word Recognition) of the Alzheimer's Disease Assessment Scale |
|  | MMSE | Total Score of Mini-Mental State Examination |
|  | RAVLT_immediate | Rey's Auditory Verbal Learning Test_Immediate Recall |
|  | RAVLT_learning | Rey's Auditory Verbal Learning Test_Learning |
|  | RAVLT_forgetting | Rey's Auditory Verbal Learning Test_Forgetting |
|  | RAVLT_perc_forgetting | Rey's Auditory Verbal Learning Test_Percent Forgetting |
|  | LDELTOTAL | Delayed total recall |
|  | TRABSCOR | Trail Making Test Part B Time |
|  | FAQ | Total Score of Functional Activities Questionnaire |
| Gene Expression | Expression values | Gene Expression Values of 120 AD patients |
| PPI | Combined Score | Combined Score from human protein physical links |

In the demographic phenotype layer, we utilized demographic and genetic features, namely age, gender, education level, race, ethnicity, and *APOE4* due to their relevance to AD risk. We applied one hot encoding on the categorical features, namely gender, race, and ethnicity. The demographic layer contains both binary and numerical values, thus the reciprocal of Gower distance ^32^ was applied to create the patient-patient similarity matrix. For neuroimaging layer and cognitive layer, we scaled each feature using min-max normalization then computed the reciprocal of Euclidean distance as patient-patient similarity value. When computing similarity scores, we ignored the features with missing values.

We created a gene network layer based on PPI. We converted STRING protein IDs to gene Entrez IDs and applied min-max normalization to the combined score. We considered the combined scores as edge weights for gene-gene interactions. The second gene network layer was constructed based on gene co-expression of 120 AD patients. The gene expression from ADNI was generated using the Affymetrix Human Genome U219 Array (Affymetrix (www.affymetrix.com), Santa Clara, CA) platform, which had 530,467 probes. We collected the normalized expression values from ADNI that were preprocessed using Robust Multi-chip Average (RMA)^33^ normalization method. To further preprocess gene expression data, we converted it from probe level to gene level by taking the mean value of probes for each corresponding gene. Then, we converted gene symbols to Entrez gene ID as the primary key for the gene networks. Reciprocal of Euclidean distance between gene expression of each gene pair was computed to generate a gene co-expression network. Then, we eliminated weakly connected edges by applying a threshold on edge weights (i.e., Reciprocal of Euclidean distance). The threshold value was chosen to ensure a balance between the number of edges and number of genes exist in gene co-expression network.

To construct the gene-patient bipartite network, we utilized the gene expression dataset. We connected a patient node to a gene node if the gene was highly upregulated in that patient. To determine which genes were highly upregulated, we first computed the median expression of each gene across all samples and considered a gene highly upregulated if its expression value was three standard deviations above the median. We used the expression values as the edge weight. Since not all patients had gene expression available, bipartite layer was only between highly expressed genes and AD patients with available expression data.

As weak edges might affect the entire network making it densely connected and causing inferior performance, we removed the weakly connected edges from the network. We selected a cutoff value (Table 2) for each network to remove the weak edges. For all the layers in the phenotype network and the PPI network layer, the cutoff value was based on the median of the edge weights. For the gene co-expression network layer, the cutoff was. selected to build a balanced network by avoiding excessive interactions at lower thresholds while preserving enough genes and connections to prevent losses at higher thresholds. Table 3 shows the final number of nodes and edges in each layer of the multiplex heterogeneous network.

**Table 2.**
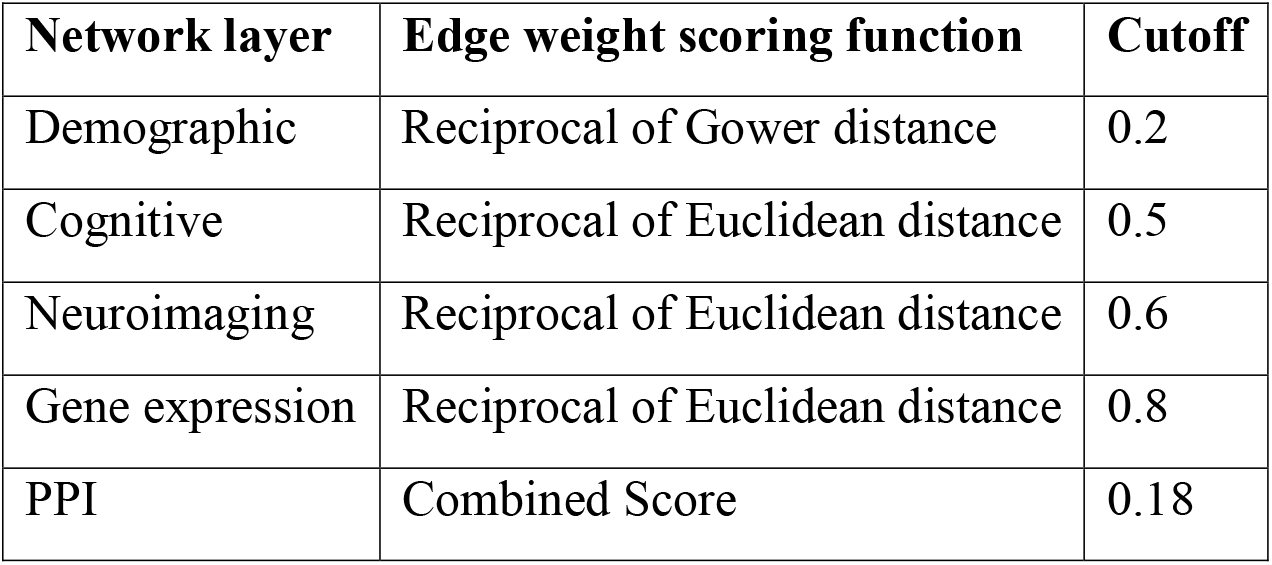
Correlation function and cutoff for each network layer.

**Table 3:** Multiplex heterogeneous network statistics.

| Network layer type | Data type | Number of edges | Number of nodes |
| --- | --- | --- | --- |
| Patient | Demographic | 146,631 | 781 |
| Patient | Cognitive | 260,858 | 805 |
| Patient | Neuroimaging | 241,891 | 795 |
| Gene | Gene expression | 3,961,187 | 12,411 |
| Gene | PPI | 405,199 | 16,872 |
| Patient-Gene | Gene expression | 9,426 | 6,816 |

### 2.2 Identifying Gold Standard AD-associated Genes

To evaluate our results, we collected a gold standard AD-associated gene set from seven different databases, namely Genetic Testing Registry (GTR)^17^ MedGen^16^ Phenotype-Genotype Integrator (PheGenI)^18^, GWAS catalog^34^, MalaCards^35^, GeneCards^36^, and NCBI Gene^37^. We searched AD in GTR database and found eight genes related to AD. In PheGenI and MedGen, we found 28 and eight genes for AD phenotype, respectively (p-value < 1e-05). We collected 153 genes from GWAS catalog selecting trait as AD and p-value < 1e-08. In GeneCards, we searched by AD relevance score greater than 30 and collected 304 genes. In MalaCards, we collected 237 genes that have ≥3 PubMed papers supporting their relevance to AD. We searched for “Alzheimer disease[All Fields] AND “Homo sapiens”[porgn] AND (“has refseqgene”[Properties] AND alive[prop])” on NCBI Gene database and found 369 AD associated RefSeq genes in human. We converted all gene symbols to Entrez gene IDs and assigned a weight to each database. We assigned a weight of 2 to GTR, MedGen, PheGenI as they are more reliable sources and a weight of 1 to GWAS, MalaCards, GeneCards and NCBI Gene databases. We computed the total score for each gene, which is the total weight of the databases the gene was reported and chose 71 genes with a score ≥ 3 (see Supplementantary Table 1).

### 2.3 Running PhenoGeneRanker

PhenoGeneRanker ^38^ is a Bioconductor R package for gene prioritization on multiplex heterogeneous networks. To run PhenoGeneRanker, we need to identify a seed gene, a gene that is known to be associated with the phenotype of interest. We also need to specify some hyperparameters to determine the influence of each component of the network.

#### 2.3.1 Seed Genes and Hyperparameters for PhenoGeneRanker

Seed nodes are the initial nodes from which the RWR algorithm starts traversing the network and returns to when “restart” occurs during the random walk. Among the gold standard AD-associated genes (see Section 2.2), we selected the genes that had a score ≥ 7 as seed genes. These genes were *APP, APOE, PSEN1, PSEN2, MPO, NOS3, PLAU, CLU, SORL1* and *ABCA7*. For each PhenoGeneRanker run, we used one of these genes as the seed. For our reference run, we used APP as the seed gene.

The hyperparameters for PhenoGeneRanker are *r*, ζ, δ, τ, η, and λ. The *r* parameter denotes the restart probability to jump back to the seed node(s). The δ and ζ parameter represents the inter-layer navigation probability for gene and phenotype networks, respectively. In other words, they control the probability of jumping to the other layers or staying on the same layer of the gene or phenotype network. For high values, the random walk will be more likely to traverse to the neighbor nodes in a different layer. The τ and φ parameter is set for assigning restart probability to each gene and phenotype network layer, respectively. These hyperparameters can be adjusted to give a higher or lower priority to restart from a particular network layer. The η parameter is used for restarting to a gene or phenotype seed in the multiplex network. A low value of η indicates that the RWR algorithm is more likely to restart on a gene seed. λ represents the probability of jumping between the gene and phenotype networks. The higher value of λ increases the dependence on the bipartite layer over gene and phenotype networks. All parameter values used in the reference run are listed in Table 4.

**Table 4.**
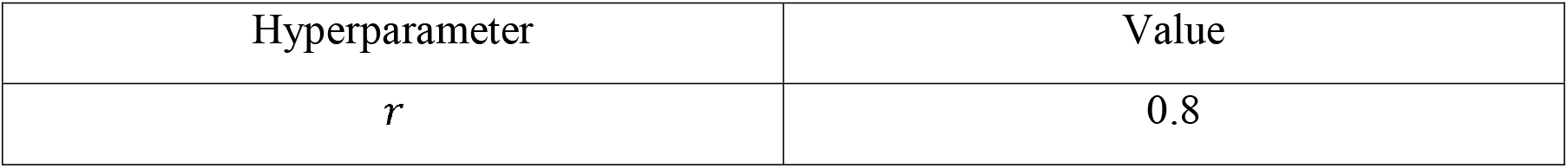
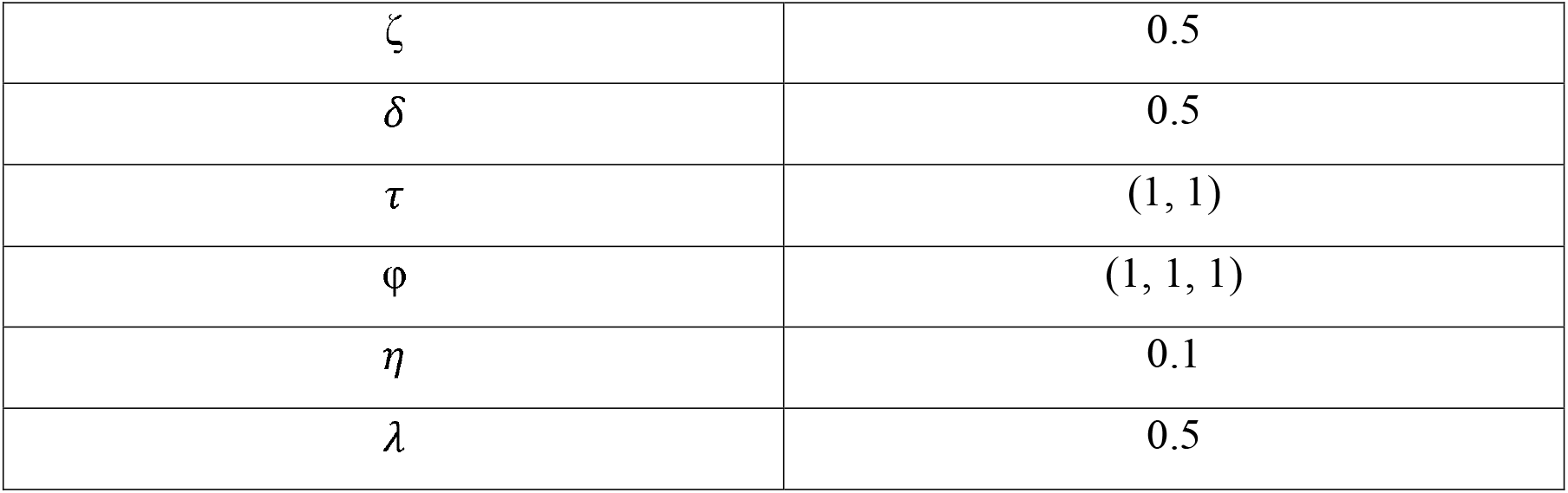
Hyperparameter values of PhenoGeneRanker used in the reference run.

| Hyperparameter | Value |
| --- | --- |
| $r$ | 0.8 |
| $\zeta$ | 0.5 |
| $\delta$ | 0.5 |
| $\tau$ | (1, 1) |
| $\varphi$ | (1, 1, 1) |
| $\eta$ | 0.1 |
| $\lambda$ | 0.5 |

## 3 Results

In this study, we built a multiplex heterogeneous network for AD using two gene layers, three phenotype layers, and one bipartite layer between phenotype and gene nodes. We ran PhenoGeneRanker on the entire multiplex heterogeneous network using APP as the seed gene (called “reference run” hereafter) to rank genes based on their association to AD. To assess the results, we performed Gene Ontology (GO) enrichment of the top 200 ranked significant genes (see Supplementary Table 2) and bottom-ranked genes. To assess the contribution of each network layer and seeds, we also ran PhenoGeneRanker with different network layer and seed gene combinations. We generated cumulative distribution function (CDF) plots to see what percentage of the gold standard AD-associated genes (see Section 2.2) were captured by PhenoGeneRanker. We also conducted literature search for the top-ranked significant genes that were not in the gold standard AD gene list. We give the details of these results in the subsequent subsections.

### 3.1 The Top-Ranked Significant Genes were Enriched in Neurodegenerative and AD related GO Terms

We performed GO enrichment for the top 200 significant genes (p-value < 0.05) using WebGestalt^39^. The results showed that the top 200 significant genes were enriched in AD-related GO biological process terms such as amyloid precursor protein metabolic process, lipid metabolic process, proteolysis, and neurogenesis ^40–43^ (False Discovery Rate (FDR) < 0.05, Table 5). These results indicate that the top genes ranked by PhenoGeneRanker were associated with AD (see Supplementary Table 3). As a negative control we repeated the same GO enrichment procedure for the bottom 200 ranked genes and found no GO terms showing that PhenoGeneRanker was able to move the irrelevant genes to the bottom of the gene prioritization list.

**Table 5:**
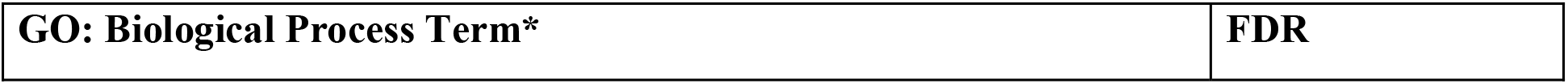
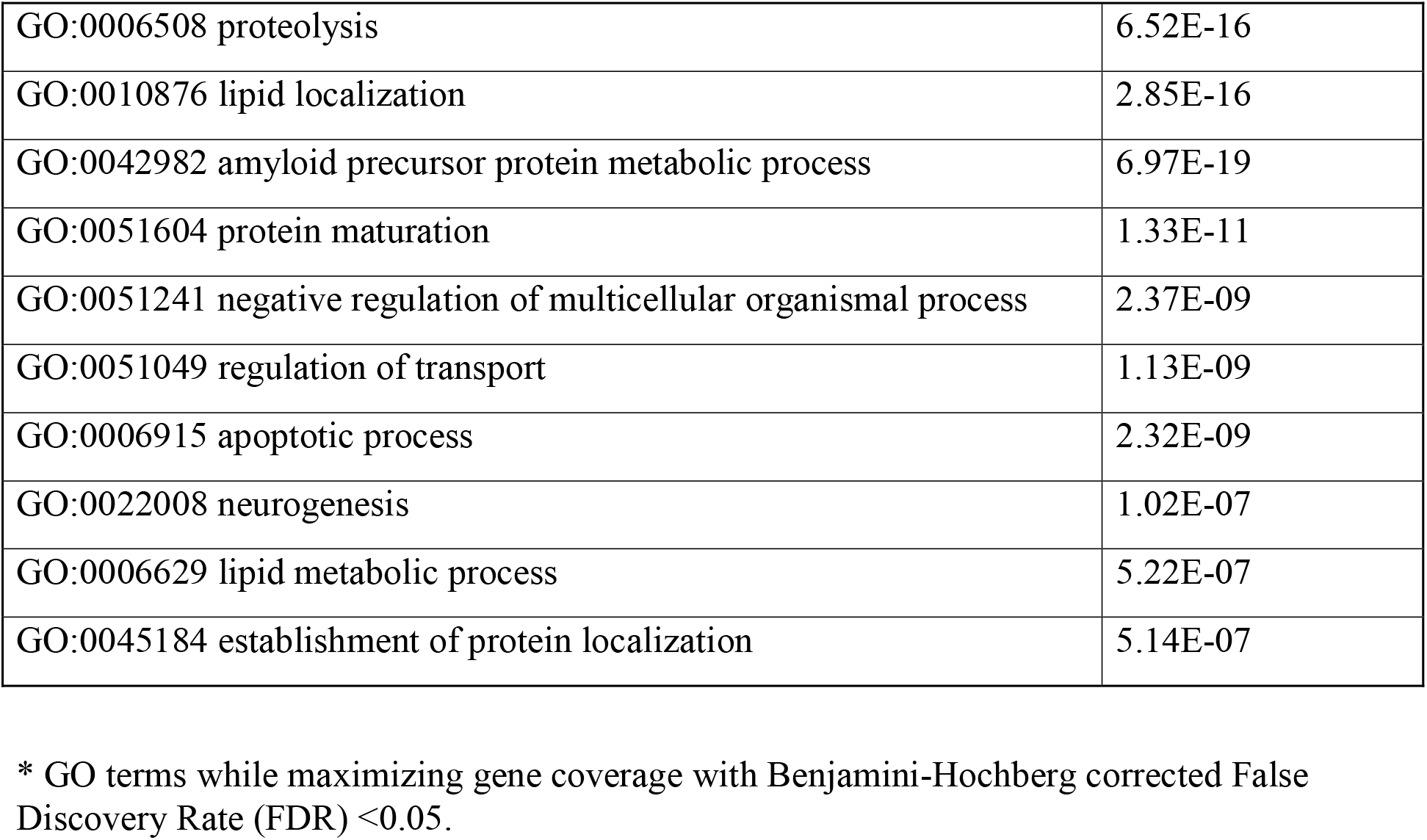
GO Biological Process enrichment of top-ranked 200 significant genes.

### 3.2 The impact of different gene layers and seed genes on the gene rankings

We compared different combinations of seed genes and network layers to check their contributions to the final gene rankings (Figure 2). Our network included 69 of 71 gold standard AD-associated genes, thus we evaluated the rankings based on these 69 genes. For each of the combinations, we kept everything same in the reference run expect for one modification. We observed that the seed gene had a high impact on the gene ranking results (Figure 2A). We checked the rankings with nine different seed genes and their impacts on the top rankings. Top 300 gene list computed using the reference run captured over 50% of genes of the 69 known AD-associated genes, whereas the coverage dropped for some other seed choices (e.g., *NOS3, PLAU*). When we ran PhenoGeneRanker using a random gene as seed gene, the coverage of the known AD genes was lowest among all seed choices. We also removed the gene co-expression layer and PPI from the networks to find their contribution in the top rankings and observed that PPI contributed higher than the gene co-expression layer (Figure 2B). We also generated a random gene network with the same number of genes and density as the original gene co-expression using Erdős–Rényi model^44^. Similarly, we generated random PPI network from original PPI network and observed the random gene layers to assess the impact of entire gene layers. We observed that random layers and using only gene co-expression layer performed poorly capturing AD-associated genes.

**Figure 2.**
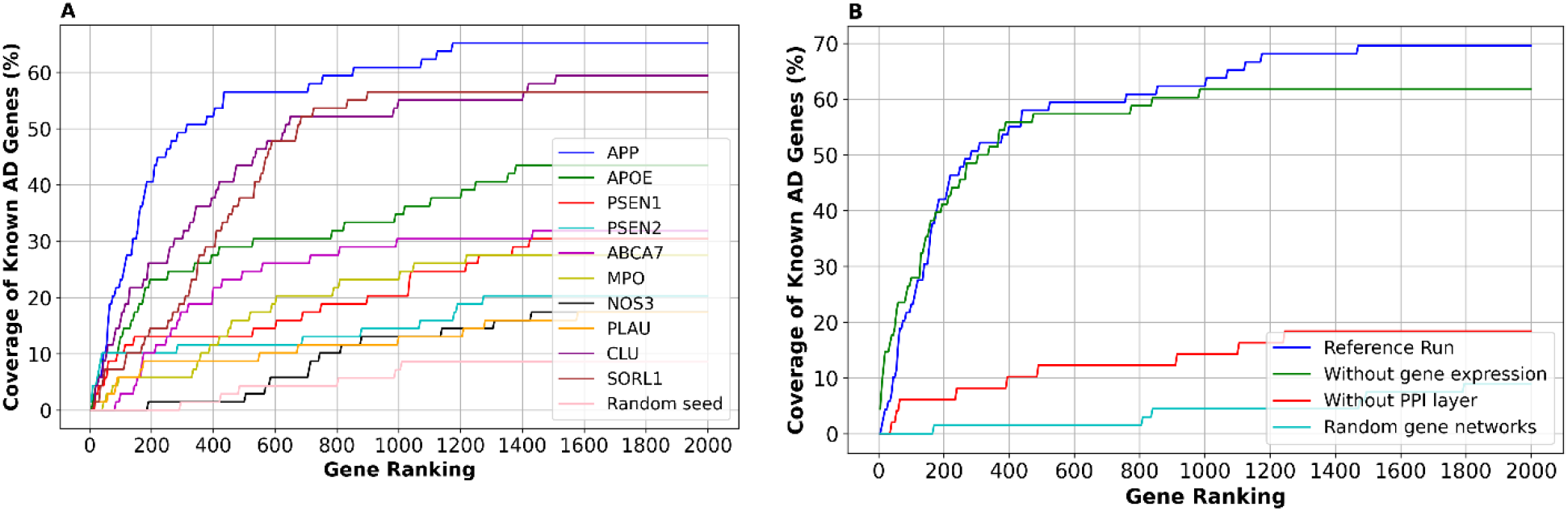
Percentage of known AD genes appearing in the top-ranked genes of PhenoGeneRanker with seed combinations and multiple networks. **A.** Percentage of known AD genes appearing in the top-ranked genes of PhenoGeneRanker with using different seed genes on the original network of the reference run. **B**. Percentage of known AD genes appearing in the top-ranked genes of PhenoGeneRanker with using different network components. The reference run uses two gene and three patient layers with APP as a seed gene. Without PPI layer, APOE is used as the seed gene because APP is not present in the gene co-expression network. Other runs are the same as the reference run except for the one change described in the figure legend. See Section 3.2 for the description of random layer generation.

### 3.3 Hyperparameter Analysis of PhenoGeneRanker

To check the effect of each PhenoGeneRanker hyperparameter on the rankings, we ran PhenoGeneRanker with multiple values for each hyperparameter for the reference run (Figure 3). We changed one hyperparameter value at a time and kept the default values for the other hyperparameters. We observed nearly identical CDF plots for different δ, ζ, and λ values (Figure 3A, 3B and 3C) indicating that the probabilities have minimal impact on the results. We observed slight differences between lower and higher *r* values, but no clear trend of improvement as *r* increases or decreases (Figure 3D). However, the CDF plots for different η and τ values were nearly identical to each other (Figure 3E and 3F).

**Figure 3.**
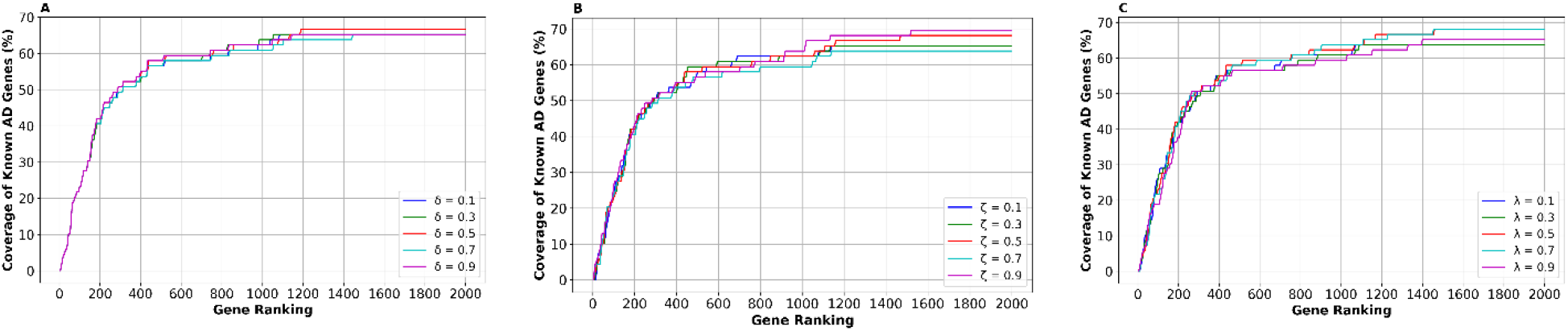
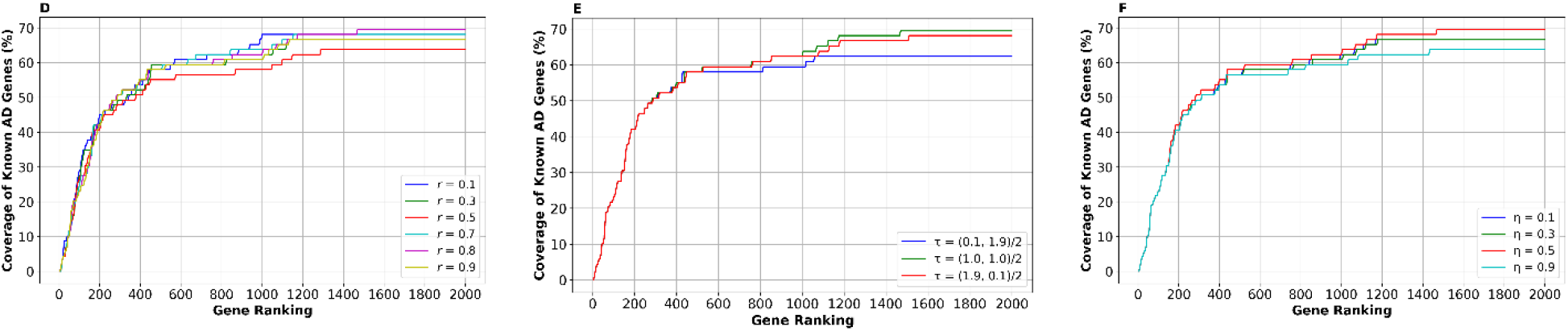
Effect of different hyperparameters on the percentage of AD gene coverage. CDF plots for different values of A) δ, B) ζ, C) λ, D) *r*, E) τ, and F) η. For each run, all the other hyperparameters were set to their default values (see Table 4) and *APP* was used as the seed gene.

### 3.4 Literature analysis of top-ranked novel AD genes

To assess if other high-ranked genes that do not appear in the list of known AD genes could be related to AD, we have selected top genes that occur in multiple runs and conducted a literature search. More specifically, we used ten different AD-related genes as a seed node (i.e., *APP PSEN1, PSEN2, APOE, MPO, PLAU ABCA7, SORL1, CLU* and *NOS3*) and ran PhenoGeneRanker to get ten different rankings. Then we selected the top common genes that appeared in top 200 significant gene lists from at least five out of ten rankings, which led to 10 genes. We removed the genes that were already in our gold standard list from the list and did literature analysis for the remaining 7 genes, which could be novel AD-related genes. We found supporting evidence for all of seven genes (Table 6). For instance, *APOA4* was reported to be decreased expression in AD patients than normal^45^, *VLDLR* slightly increased in AD conditions^46^.

**Table 6.**
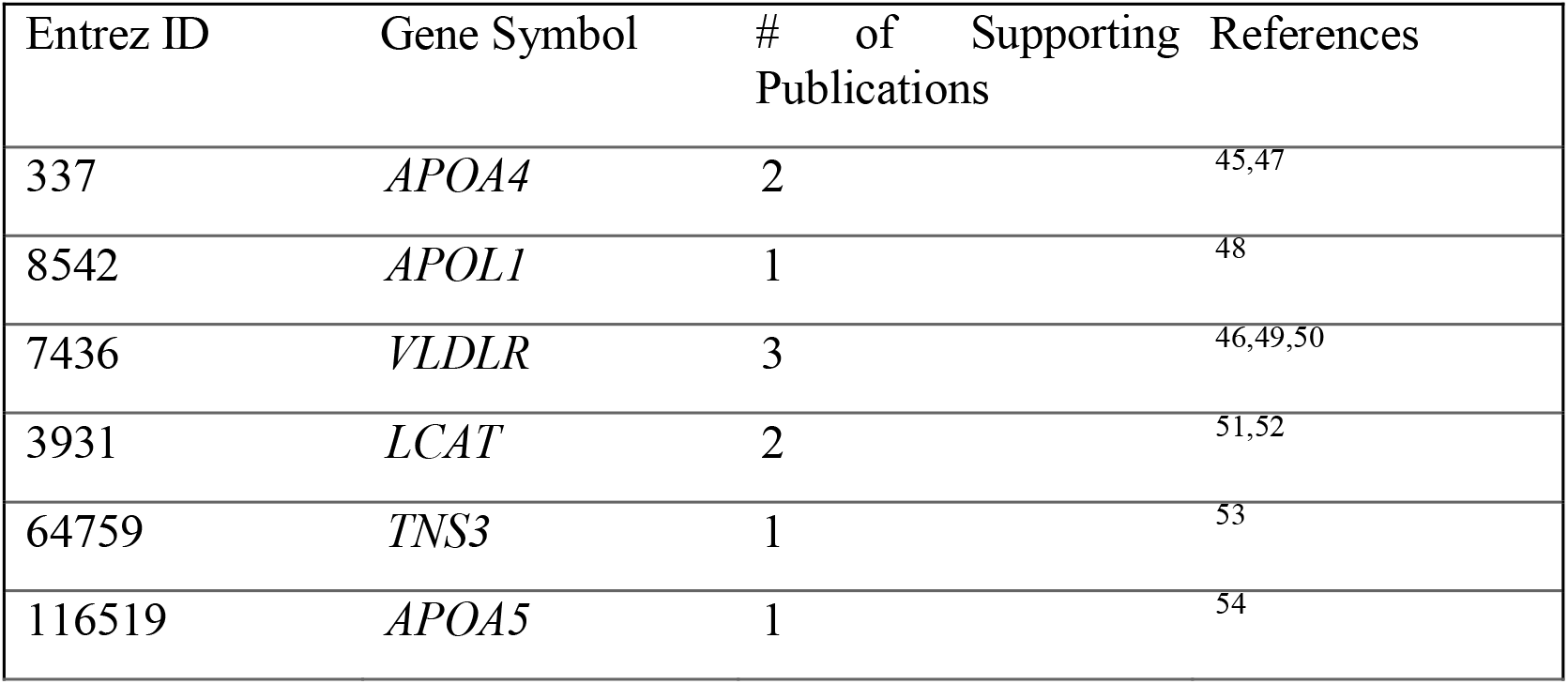
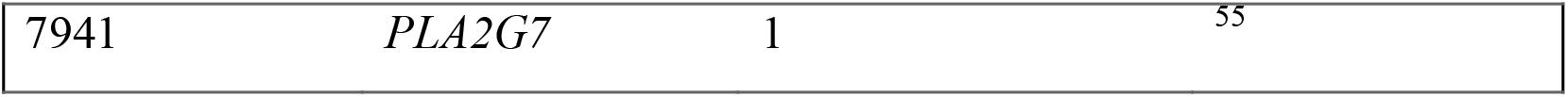
Novel AD genes, appeared in multiple rankings of PhenoGeneRanker using different seed genes, with number of supporting publications and their references.

## 4 Discussion

In this study, we constructed a multiplex heterogeneous network for AD that consists of two gene layers, three phenotype layers, and a bipartite layer connecting patient phenotype nodes to gene nodes. In the phenotype layer, each node represents an AD patient, the links with the other AD patients were based on their similarity scores calculated using demographic, cognitive, and neuroimaging data obtained from ADNI. For the gene layer, each node represents a gene, and connections between the two genes were based on PPI and gene expression values. After building complete multiplex heterogeneous network, the PhenoGeneRanker tool was executed on the network to rank genes by their association with AD known genes. Finally, potential AD-related genes were prioritized through the analysis of multimodal AD associated datasets.

We performed GO enrichment analysis for the top 200 and bottom 200 ranked genes to observe whether overrepresented biological processes were related to AD. Our top 200 significant genes were enriched in GO biological process terms such as *APP* metabolic process, lipid metabolic process, proteolysis, apoptotic process and neurogenesis. Improper metabolism of *APP* leads to the accumulation of amyloid-beta plaques, and abnormal apoptosis leads to the loss of neurons in the brain areas like the cerebral cortex and hippocampus^56,57^. However, the bottom 200 ranked genes were found to have no significant GO terms which indicates that the PhenoGeneRanker tool effectively pushed irrelevant genes to the bottom.

We checked the impact of different gene layers and seed genes on the gene rankings. We observed that the PPI layer contributed more than the gene expression layer. We assumed that it was due to the small sample size of gene expression data. Although our phenotype layers included nearly 800 AD patients, the gene expression layer had available data from only 120 AD patients. We selected well-known AD-associated genes as seed genes to evaluate their effectiveness in capturing gold-standard AD-associated genes. Among them, our reference run captured over 50% of these genes within the top 300 significant ranked genes. This indicates that our proposed approach was able to prioritize relevant genes in the top list. However, there is a room for improvement to be able to capture additional genes that were not ranked in the top listed genes. We also noticed the PhenoGeneRanker’s robustness to changes in hyperparameters such as δ, ζ, and λ (Figure 3). The model’s performance remained consistent which indicated its stability and reliability across different parameter settings.

We generated 10 different gene rankings using ten different AD-known genes as a seed: *APP, PSEN1, PSEN2, APOE, MPO, PLAU, ABCA7, SORL1, CLU*, and *NOS3*, and selected the genes if they appeared in the top 200 gene list in at least five out of the ten rankings. We found seven genes that are not listed in our gold-standard AD-associated genes. The identified seven genes were consistently ranked highly across different seed-based analyses which increased the confidence that these genes could be relevant to AD. We performed literature search for these genes to examine their potential relevance to AD biology. We found out that reduced levels of *APOA4* are linked to a higher risk of developing AD^45^. Additionally, increased levels of *APOL1* have been observed in the frontal cortex of individuals with frontotemporal lobar degeneration (FTLD) that leads to dementia^48^. Polymorphisms in the *VLDLR* gene may be associated with AD in Japanese individuals, particularly those with an earlier age of onset^46^. Furthermore, increased levels of *PLA2G7* are linked to a risk factor for developing AD^55^, while CSF analysis has shown reduced *LCAT* activity in the plasma lipoprotein profiles of AD patients compared to controls^51^. Although *TNS3* is not directly associated with AD, it interacts with *APOE* and *CLU*, which have both been linked to AD^53^. Lastly, genetic vulnerability related to *APOA5* may influence the association between triglyceride levels and the risk of AD in women where low triglyceride levels are linked to a decreased risk of AD^54^.

In our current study, we used gene expression and PPI networks; however, in future work, we plan to expand our gene layers by integrating DNA methylation and SNP data. Additionally, we only considered the connections between genes that might cause the loss of some important information about the genes by not considering the values coming from AD patients. In the future, our network will be generated based on gene connection and their values coming from different data modalities. Another limitation of our study is that the data predominantly represents a white population (751 out of 805 participants). Utilizing datasets from more diverse populations would be useful to identify additional genes that may be associated with AD in different demographic groups. To address this issue, we aim to select racially balanced datasets in future analyses.

## 5. Conclusion

In this study, we generated a multiplex heterogeneous network for AD and applied PhenoGeneRanker on this network to identify putative AD-related genes. We used *APP* as seed gene because of having strong relationships with AD from several studies. Our results showed that the top-ranked genes were enriched in AD-related GO terms. Several known AD genes appear at the top of the ranking. Our top 200 gene list captured about 40% of genes that were reported as AD-associated. We also found some genes that were ranked highly in multiple PhenoGeneRanker runs, which had supporting literature for their potential association to AD. The integration of multimodal datasets for AD using multiplex heterogeneous network and PhenoGeneRanker tool may be utilized for finding disease-associated genes.

## Acknowledgements

Data collection and sharing for this project was funded by the Alzheimer’s Disease Neuroimaging Initiative (ADNI) (National Institutes of Health Grant U01 AG024904) and DOD ADNI (Department of Defense award number W81XWH-12-2-0012). ADNI is funded by the National Institute on Aging, the National Institute of Biomedical Imaging and Bioengineering, and through generous contributions from the following: AbbVie, Alzheimer’s Association; Alzheimer’s Drug Discovery Foundation; Araclon Biotech; BioClinica, Inc.; Biogen; Bristol-Myers Squibb Company; CereSpir, Inc.; Cogstate; Eisai Inc.; Elan Pharmaceuticals, Inc.; Eli Lilly and Company; EuroImmun; F. Hoffmann-La Roche Ltd and its affiliated company Genentech, Inc.; Fujirebio; GE Healthcare; IXICO Ltd.; Janssen Alzheimer Immunotherapy Research & Development, LLC.; Johnson & Johnson Pharmaceutical Research & Development LLC.; Lumosity; Lundbeck; Merck & Co., Inc.; Meso Scale Diagnostics, LLC.; NeuroRx Research; Neurotrack Technologies; Novartis Pharmaceuticals Corporation; Pfizer Inc.; Piramal Imaging; Servier; Takeda Pharmaceutical Company; and Transition Therapeutics. The Canadian Institutes of Health Research is providing funds to support ADNI clinical sites in Canada. Private sector contributions are facilitated by the Foundation for the National Institutes of Health (www.fnih.org). The grantee organization is the Northern California Institute for Research and Education, and the study is coordinated by the Alzheimer’s Therapeutic Research Institute at the University of Southern California. ADNI data are disseminated by the Laboratory for Neuro Imaging at the University of Southern California.

## Consent Statement

The ADNI obtained written informed consent from all participants.

## References

1. Patterson, C. (2018). World alzheimer report 2018.

2. Pudelewicz, A., Talarska, D. & Bączyk, G. Burden of caregivers of patients with Alzheimer’s disease. Scand. J. Caring Sci. 33, 336–341 (2019).

3. Jayadev, S. et al. Alzheimer’s disease phenotypes and genotypes associated with mutations in presenilin 2. Brain 133, 1143–1154 (2010).

4. Bloom, G. S. Amyloid-β and Tau: The Trigger and Bullet in Alzheimer Disease Pathogenesis. JAMA Neurol. 71, 505 (2014).

5. Gulisano, W. et al. Role of Amyloid-β and Tau Proteins in Alzheimer’s Disease: Confuting the Amyloid Cascade. J. Alzheimers Dis. 64, S611–S631 (2018).

6. Lloret et al. When Does Alzheimer′s Disease Really Start? The Role of Biomarkers. Int. J. Mol. Sci. 20, 5536 (2019).

7. Palmqvist, S. et al. Detailed comparison of amyloid PET and CSF biomarkers for identifying early Alzheimer disease. Neurology 85, 1240–1249 (2015).

8. Ding, J. et al. Tau-PET abnormality as a biomarker for Alzheimer’s disease staging and early detection: a topological perspective. Cereb. Cortex 33, 10649–10659 (2023).

9. Hu, Z., Wang, Z., Jin, Y. & Hou, W. VGG-TSwinformer: Transformer-based deep learning model for early Alzheimer’s disease prediction. Comput. Methods Programs Biomed. 229, 107291 (2023).

10. Al Olaimat, M., Martinez, J., Saeed, F., Bozdag, S., & Alzheimer’s Disease Neuroimaging Initiative. PPAD: a deep learning architecture to predict progression of Alzheimer’s disease. Bioinformatics 39, i149–i157 (2023).

11. Olaimat, M. A. & Bozdag, S. TA-RNN: an Attention-based Time-aware Recurrent Neural Network Architecture for Electronic Health Records. Preprint at http://arxiv.org/abs/2401.14694 (2024).

12. Akhavan Aghdam, M., Bozdag, S., Saeed, F., & Alzheimer’s Disease Neuroimaging Initiative. PVTAD: ALZHEIMER’S DISEASE DIAGNOSIS USING PYRAMID VISION TRANSFORMER APPLIED TO WHITE MATTER OF T1-WEIGHTED STRUCTURAL MRI DATA. Preprint at 10.1101/2023.11.17.567617 (2023).

13. Harold, D. et al. Genome-wide association study identifies variants at CLU and PICALM associated with Alzheimer’s disease. Nat. Genet. 41, 1088–1093 (2009).

14. the Alzheimer’s Disease Neuroimaging Initiative et al. Common variants at ABCA7, MS4A6A/MS4A4E, EPHA1, CD33 and CD2AP are associated with Alzheimer’s disease. Nat. Genet. 43, 429–435 (2011).

15. Naj, A. C. et al. Common variants at MS4A4/MS4A6E, CD2AP, CD33 and EPHA1 are associated with late-onset Alzheimer’s disease. Nat. Genet. 43, 436–441 (2011).

16. Database resources of the National Center for Biotechnology Information. Nucleic Acids Res. 44, D7–D19 (2016).

17. Sayers, E. W. et al. Database resources of the National Center for Biotechnology Information in 2023. Nucleic Acids Res. 51, D29–D38 (2023).

18. Ramos, E. M. et al. Phenotype–Genotype Integrator (PheGenI): synthesizing genome-wide association study (GWAS) data with existing genomic resources. Eur. J. Hum. Genet. 22, 144–147 (2014).

19. Bertram, L., McQueen, M. B., Mullin, K., Blacker, D. & Tanzi, R. E. Systematic meta-analyses of Alzheimer disease genetic association studies: the AlzGene database. Nat. Genet. 39, 17–23 (2007).

20. Reynolds, W. F. et al. MPO and APOEε4 polymorphisms interact to increase risk for AD in Finnish males. Neurology 55, 1284–1290 (2000).

21. Zhao, Q.-F., Yu, J.-T.Tan, M.-S. & Tan, L. ABCA7 in Alzheimer’s Disease. Mol. Neurobiol. 51, 1008–1016 (2015).

22. Šerý, O., Povová, J., Míšek, I., Pešák, L. & Janout, V. Molecular mechanisms of neuropathological changes in Alzheimer’s disease: a review. Folia Neuropathol. 1, 1–9 (2013).

23. Xia, L.-Y., Tang, L., Huang, H. & Luo, J. Identification of Potential Driver Genes and Pathways Based on Transcriptomics Data in Alzheimer’s Disease. Front. Aging Neurosci. 14, 752858 (2022).

24. Liao, W. et al. Identification of candidate genes associated with clinical onset of Alzheimer’s disease. Front. Neurosci. 16, 1060111 (2022).

25. Cowen, L., Ideker, T., Raphael, B. J. & Sharan, R. Network propagation: a universal amplifier of genetic associations. Nat. Rev. Genet. 18, 551–562 (2017).

26. Erten, S., Bebek, G., Ewing, R. M. & Koyutürk, M. DA DA: Degree-Aware Algorithms for Network-Based Disease Gene Prioritization. BioData Min. 4, 19 (2011).

27. Köhler, S., Bauer, S., Horn, D. & Robinson, P. N. Walking the Interactome for Prioritization of Candidate Disease Genes. Am. J. Hum. Genet. 82, 949–958 (2008).

28. Valdeolivas, A. et al. Random walk with restart on multiplex and heterogeneous biological networks. Bioinformatics 35, 497–505 (2019).

29. Dursun, C., Kwitek, A. E. & Bozdag, S. PhenoGeneRanker: Gene and Phenotype Prioritization Using Multiplex Heterogeneous Networks. IEEE/ACM Trans. Comput. Biol. Bioinform. 19, 2950–2962 (2022).

30. Alzheimer’s Disease Neuroimaging Initiative.(2023). ADNIMERGE: Alzheimer’s Disease Neuroimaging Initiative. R package version 0.0.1. Available online: https://adni.bitbucket.io.

31. Szklarczyk, D. et al. The STRING database in 2023: protein–protein association networks and functional enrichment analyses for any sequenced genome of interest. Nucleic Acids Res. 51, D638–D646 (2023).

32. Gower, J. C. A General Coefficient of Similarity and Some of Its Properties. Biometrics 27, 857 (1971).

33. Irizarry, R. A. Exploration, normalization, and summaries of high density oligonucleotide array probe level data. Biostatistics 4, 249–264 (2003).

34. Buniello, A. et al. The NHGRI-EBI GWAS Catalog of published genome-wide association studies, targeted arrays and summary statistics 2019. Nucleic Acids Res. 47, D1005–D1012 (2019).

35. Rappaport, N. et al. MalaCards: an amalgamated human disease compendium with diverse clinical and genetic annotation and structured search. Nucleic Acids Res. 45, D877–D887 (2017).

36. Stelzer, G. et al. The GeneCards Suite: From Gene Data Mining to Disease Genome Sequence Analyses. Curr. Protoc. Bioinforma. 54, (2016).

37. Brown, G. R. et al. Gene: a gene-centered information resource at NCBI. Nucleic Acids Res. 43, D36–D42 (2015).

38. Cagatay Dursun <cagataydursun@gmail. com> [aut cre]. PhenoGeneRanker. Bioconductor 10.18129/B9.BIOC.PHENOGENERANKER.

39. Liao, Y., Wang, J., Jaehnig, E. J., Shi, Z. & Zhang, B. WebGestalt 2019: gene set analysis toolkit with revamped UIs and APIs. Nucleic Acids Res. 47, W199–W205 (2019).

40. Ma, G. et al. Differential Expression of mRNAs in the Brain Tissues of Patients with Alzheimer’s Disease Based on GEO Expression Profile and Its Clinical Significance. BioMed Res. Int. 2019, 1–9 (2019).

41. Fan, Z. et al. Identification of Potential Therapeutic Targets of Alzheimer’s Disease By Weighted Gene Co-Expression Network Analysis. Chin. Med. Sci. J. 35, 330 (2020).

42. Sun, T. et al. Comprehensive analysis of dysregulated circular RNAs and construction of a ceRNA network involved in the pathology of Alzheimer’s disease in a 5 × FAD mouse model. Front. Aging Neurosci. 14, 1020699 (2022).

43. Sung, P.-S., Lin, P.-Y., Liu, C.-H.Su, H.-C. & Tsai, K.-J. Neuroinflammation and Neurogenesis in Alzheimer’s Disease and Potential Therapeutic Approaches. Int. J. Mol. Sci. 21, 701 (2020).

44. Erdős, P. & Rényi, A. On random graphs. I. Publ. Math. Debr. 6, 290–297 (2022).

45. Lin, Q., Cao, Y. & Gao, J. Decreased expression of the APOA1–APOC3–APOA4 gene cluster is associated with risk of Alzheimer’s disease. Drug Des. Devel. Ther. 5421 (2015) doi:10.2147/DDDT.S89279.

46. Yamanaka, H. et al. Genetic Risk Factors in Japanese Alzheimer’s Disease Patients: α1-ACT, VLDLR, and ApoE. Neurobiol. Aging 19, S43–S46 (1998).

47. Nicolaides, N. C. et al. Plasma Proteomics in Healthy Subjects with Differences in Tissue Glucocorticoid Sensitivity Identifies A Novel Proteomic Signature. Biomedicines 10, 184 (2022).

48. Hok-A-Hin, Y. S. et al. Apolipoprotein L1 is increased in frontotemporal lobar degeneration post-mortem brain but not in ante-mortem cerebrospinal fluid. Neurobiol. Dis. 172, 105813 (2022).

49. Holtzman, D. M., Herz, J. & Bu, G. Apolipoprotein E and Apolipoprotein E Receptors: Normal Biology and Roles in Alzheimer Disease. Cold Spring Harb. Perspect. Med. 2, a006312–a006312 (2012).

50. Lane-Donovan, C. & Herz, J. The ApoE receptors Vldlr and Apoer2 in central nervous system function and disease. J. Lipid Res. 58, 1036–1043 (2017).

51. Turri, M. et al. Plasma and cerebrospinal fluid cholesterol esterification is hampered in Alzheimer’s disease. Alzheimers Res. Ther. 15, 95 (2023).

52. Hong, B. V. et al. High-Density Lipoprotein Changes in Alzheimer’s Disease Are APOE Genotype-Specific. Biomedicines 10, 1495 (2022).

53. Huang, Y. et al. A machine learning approach to brain epigenetic analysis reveals kinases associated with Alzheimer’s disease. Nat. Commun. 12, 4472 (2021).

54. Ancelin, M.-L. et al. Sex Differences in the Associations Between Lipid Levels and Incident Dementia. J. Alzheimers Dis. 34, 519–528 (2013).

55. Liu, Q. et al. Structural and Thermodynamic Characterization of Protein–Ligand Interactions Formed between Lipoprotein-Associated Phospholipase A2 and Inhibitors. J. Med. Chem. 59, 5115–5120 (2016).

56. Zheng, H. & Koo, E. H. Biology and pathophysiology of the amyloid precursor protein. Mol. Neurodegener. 6, 27 (2011).

57. Sharma, V. K., Singh, T. G., Singh, S., Garg, N. & Dhiman, S. Apoptotic Pathways and Alzheimer’s Disease: Probing Therapeutic Potential. Neurochem. Res. 46, 3103–3122 (2021).

