## Supplementary Table 1 for "Identifying Alzheimer’s disease-associated genes using PhenoGeneRanker"

**Supplementary Table 1:** AD-associated genes across seven databases. + means the gene was reported in that database. The number in parentheses denote the weight of each database. We selected the genes that had a total score ≥ 3 as the gold standard AD-associated genes.

| Gene Symbol | GTR (2) | MedGen (2) | PheGenI (2) | GWAS (1) | MalaCards (1) | GeneCards (1) | NCBI gene (1) |
| --- | --- | --- | --- | --- | --- | --- | --- |
| APOE | + | + | + | + | + | + | + |
| ABCA7 | + | + | + | + |  | + | + |
| APP | + | + |  | + | + | + | + |
| MPO | + | + |  |  | + | + | + |
| NOS3 | + | + |  |  | + | + | + |
| PLAU | + | + |  |  | + | + | + |
| PSEN1 | + | + |  |  | + | + | + |
| PSEN2 | + | + |  |  | + | + | + |
| CLU |  |  | + | + | + | + | + |
| SORL1 |  |  | + | + | + | + | + |
| APOC1 |  |  | + | + | + |  | + |
| CR1 |  |  | + | + |  | + | + |
| TOMM40 |  |  | + | + |  | + | + |
| ADAM10 | + |  |  |  | + | + | + |
| MAPT | + |  |  |  | + | + | + |
| SNCA | + |  |  |  | + | + | + |
| ABCA1 |  |  |  | + | + | + | + |
| BIN1 |  |  | + | + |  |  | + |
| GRN |  |  |  | + | + | + | + |
| MME |  |  |  | + | + | + | + |
| MS4A6A |  |  | + | + |  |  | + |
| NECTIN2 |  |  | + | + |  |  | + |
| PICALM |  |  | + | + |  |  | + |
| GBA1 | + |  |  |  | + | + |  |
| ITM2B | + |  |  |  | + |  | + |
| SNCB | + |  |  |  |  | + | + |
| ADAM17 |  |  |  | + |  | + | + |
| CTSB |  |  |  | + | + |  | + |
| EPHA1-AS1 |  |  | + | + |  |  |  |
| MS4A2 |  |  | + | + |  |  |  |
| NTF3 |  |  |  | + | + |  | + |
| PTK2B |  |  | + | + |  |  |  |
| TREM2 |  |  |  | + |  | + | + |
| A2M |  |  |  |  | + | + | + |
| ACHE |  |  |  |  | + | + | + |
| AGER |  |  |  |  | + | + | + |
| APOB |  |  |  |  | + | + | + |
| BACE1 |  |  |  |  | + | + | + |
| BCHE |  |  |  |  | + | + | + |
| BDNF |  |  |  |  | + | + | + |
| BLMH |  |  |  |  | + | + | + |
| CBS |  |  |  |  | + | + | + |
| CDK5 |  |  |  |  | + | + | + |
| CHAT |  |  |  |  | + | + | + |
| COMT |  |  |  |  | + | + | + |
| CST3 |  |  |  |  | + | + | + |
| CTSD |  |  |  |  | + | + | + |
| CYP46A1 |  |  |  |  | + | + | + |
| GSK3B |  |  |  |  | + | + | + |
| HFE |  |  |  |  | + | + | + |
| HMOX1 |  |  |  |  | + | + | + |
| HSD17B10 |  |  |  |  | + | + | + |
| IDE |  |  |  |  | + | + | + |
| IL1A |  |  |  |  | + | + | + |
| IL1B |  |  |  |  | + | + | + |
| LRP1 |  |  |  |  | + | + | + |
| LRRK2 |  |  |  |  | + | + | + |
| NCSTN |  |  |  |  | + | + | + |
| PON1 |  |  |  |  | + | + | + |
| PPARG |  |  |  |  | + | + | + |
| PRKN |  |  |  |  | + | + | + |
| PRNP |  |  |  |  | + | + | + |
| PSENEN |  |  |  |  | + | + | + |
| PTGS2 |  |  |  |  | + | + | + |
| SERPINA3 |  |  |  |  | + | + | + |
| SLC6A3 |  |  |  |  | + | + | + |
| SOD1 |  |  |  |  | + | + | + |
| SQSTM1 |  |  |  |  | + | + | + |
| TARDBP |  |  |  |  | + | + | + |
| TTR |  |  |  |  | + | + | + |
| UCHL1 |  |  |  |  | + | + | + |
