## Supplementary Table 2 for "Identifying Alzheimer’s disease-associated genes using PhenoGeneRanker"

**Supplementary Table 2:** AD-associated top 200 significant genes ranked by PhenoGeneRanker.

| **Gene** | **Score** | **P_value** |
| --- | --- | --- |
| 10531 | 0.000112089 | 0 |
| 5653 | 0.000111186 | 0 |
| 8883 | 8.08E-05 | 0.006006006 |
| 27242 | 7.87E-05 | 0.003003003 |
| 152330 | 7.86E-05 | 0 |
| 10307 | 7.85E-05 | 0.012012012 |
| 9546 | 7.79E-05 | 0 |
| 143458 | 7.16E-05 | 0 |
| 320 | 6.50E-05 | 0.006006006 |
| 6653 | 6.41E-05 | 0.015015015 |
| 83941 | 6.40E-05 | 0 |
| 54103 | 6.32E-05 | 0 |
| 23385 | 6.23E-05 | 0 |
| 9911 | 6.12E-05 | 0.024024024 |
| 26020 | 6.05E-05 | 0 |
| 321 | 5.96E-05 | 0 |
| 51454 | 5.49E-05 | 0.003003003 |
| 43849 | 5.47E-05 | 0 |
| 23621 | 5.34E-05 | 0.003003003 |
| 5067 | 5.23E-05 | 0 |
| 10418 | 5.21E-05 | 0 |
| 323 | 4.95E-05 | 0 |
| 51734 | 4.82E-05 | 0 |
| 200576 | 4.79E-05 | 0.012012012 |
| 53353 | 4.76E-05 | 0.009009009 |
| 6900 | 4.73E-05 | 0.006006006 |
| 6440 | 4.65E-05 | 0.012012012 |
| 7092 | 4.64E-05 | 0 |
| 9445 | 4.57E-05 | 0.003003003 |
| 57136 | 4.48E-05 | 0 |
| 4038 | 4.48E-05 | 0.018018018 |
| 414325 | 4.42E-05 | 0 |
| 2358 | 4.40E-05 | 0 |
| 30061 | 4.39E-05 | 0 |
| 51107 | 4.26E-05 | 0.006006006 |
| 124906744 | 4.23E-05 | 0 |
| 55894 | 4.23E-05 | 0 |
| 3934 | 4.21E-05 | 0 |
| 5663 | 4.07E-05 | 0.009009009 |
| 2931 | 4.02E-05 | 0.003003003 |
| 43 | 4.01E-05 | 0.018018018 |
| 3416 | 4.01E-05 | 0.024024024 |
| 10513 | 3.79E-05 | 0.003003003 |
| 336 | 3.68E-05 | 0.033033033 |
| 2213 | 3.65E-05 | 0.012012012 |
| 2160 | 3.64E-05 | 0.009009009 |
| 4763 | 3.63E-05 | 0.021021021 |
| 322 | 3.63E-05 | 0.009009009 |
| 3931 | 3.60E-05 | 0.006006006 |
| 9039 | 3.60E-05 | 0.006006006 |
| 22883 | 3.59E-05 | 0.003003003 |
| 56899 | 3.58E-05 | 0 |
| 4035 | 3.55E-05 | 0.015015015 |
| 839 | 3.54E-05 | 0.006006006 |
| 151056 | 3.48E-05 | 0.003003003 |
| 337 | 3.48E-05 | 0.009009009 |
| 7276 | 3.48E-05 | 0.042042042 |
| 7941 | 3.39E-05 | 0.006006006 |
| 1471 | 3.34E-05 | 0.009009009 |
| 642 | 3.31E-05 | 0.006006006 |
| 81890 | 3.30E-05 | 0 |
| 5621 | 3.29E-05 | 0.006006006 |
| 334 | 3.26E-05 | 0.003003003 |
| 1191 | 3.24E-05 | 0.012012012 |
| 4804 | 3.22E-05 | 0.018018018 |
| 350 | 3.22E-05 | 0.015015015 |
| 3375 | 3.19E-05 | 0.012012012 |
| 345 | 3.18E-05 | 0.018018018 |
| 3329 | 3.14E-05 | 0.03003003 |
| 8504 | 3.14E-05 | 0.006006006 |
| 26085 | 3.12E-05 | 0.003003003 |
| 9479 | 3.06E-05 | 0.024024024 |
| 3028 | 3.05E-05 | 0.018018018 |
| 5481 | 3.02E-05 | 0.012048193 |
| 43847 | 3.00E-05 | 0.015015015 |
| 6291 | 2.99E-05 | 0.006024096 |
| 346 | 2.94E-05 | 0.006006006 |
| 51449 | 2.92E-05 | 0.006006006 |
| 116519 | 2.86E-05 | 0.021021021 |
| 6855 | 2.86E-05 | 0.006006006 |
| 55697 | 2.85E-05 | 0.024024024 |
| 1071 | 2.85E-05 | 0.042042042 |
| 55851 | 2.83E-05 | 0.015015015 |
| 129521 | 2.82E-05 | 0.018018018 |
| 1600 | 2.81E-05 | 0.033033033 |
| 7804 | 2.79E-05 | 0.024024024 |
| 5646 | 2.77E-05 | 0.018018018 |
| 3163 | 2.77E-05 | 0.033033033 |
| 10288 | 2.77E-05 | 0 |
| 55937 | 2.75E-05 | 0.036036036 |
| 53942 | 2.70E-05 | 0.003003003 |
| 11012 | 2.70E-05 | 0 |
| 8542 | 2.69E-05 | 0.039039039 |
| 1622 | 2.69E-05 | 0.009009009 |
| 64759 | 2.66E-05 | 0.006006006 |
| 341 | 2.64E-05 | 0.012012012 |
| 4018 | 2.62E-05 | 0.012012012 |
| 7917 | 2.60E-05 | 0.015015015 |
| 9179 | 2.56E-05 | 0.006006006 |
| 5118 | 2.56E-05 | 0.006006006 |
| 1139 | 2.54E-05 | 0.003003003 |
| 2923 | 2.53E-05 | 0.018018018 |
| 51608 | 2.51E-05 | 0.021021021 |
| 65078 | 2.51E-05 | 0.021021021 |
| 5654 | 2.50E-05 | 0 |
| 11202 | 2.49E-05 | 0.003003003 |
| 3069 | 2.48E-05 | 0.039039039 |
| 5649 | 2.47E-05 | 0.012012012 |
| 26060 | 2.46E-05 | 0.012012012 |
| 1814 | 2.46E-05 | 0.03003003 |
| 344 | 2.42E-05 | 0.006006006 |
| 2771 | 2.42E-05 | 0.018018018 |
| 4924 | 2.40E-05 | 0.015015015 |
| 5444 | 2.38E-05 | 0.024024024 |
| 4601 | 2.38E-05 | 0.009009009 |
| 55754 | 2.35E-05 | 0.012012012 |
| 348 | 2.35E-05 | 0.045045045 |
| 5645 | 2.32E-05 | 0.003003003 |
| 55911 | 2.32E-05 | 0.009009009 |
| 55554 | 2.30E-05 | 0.003003003 |
| 29979 | 2.28E-05 | 0.015015015 |
| 335 | 2.28E-05 | 0.048048048 |
| 11018 | 2.24E-05 | 0.039039039 |
| 100528017 | 2.23E-05 | 0.012012012 |
| 55829 | 2.22E-05 | 0.015015015 |
| 51466 | 2.22E-05 | 0.012012012 |
| 1719 | 2.19E-05 | 0.015015015 |
| 57216 | 2.19E-05 | 0.015015015 |
| 5367 | 2.19E-05 | 0.024024024 |
| 51599 | 2.17E-05 | 0.006006006 |
| 27255 | 2.15E-05 | 0.009009009 |
| 79135 | 2.15E-05 | 0.012012012 |
| 972 | 2.14E-05 | 0.033033033 |
| 10439 | 2.14E-05 | 0.012012012 |
| 10452 | 2.10E-05 | 0.03003003 |
| 10874 | 2.08E-05 | 0.015015015 |
| 6620 | 2.07E-05 | 0.009009009 |
| 1410 | 2.06E-05 | 0.012012012 |
| 6750 | 2.06E-05 | 0.03003003 |
| 8650 | 2.05E-05 | 0.006006006 |
| 6449 | 2.04E-05 | 0.018018018 |
| 2903 | 2.03E-05 | 0.042042042 |
| 10524 | 2.01E-05 | 0.039039039 |
| 5644 | 2.00E-05 | 0.018018018 |
| 6289 | 2.00E-05 | 0.012012012 |
| 5360 | 1.98E-05 | 0.033033033 |
| 10498 | 1.97E-05 | 0.006006006 |
| 6368 | 1.95E-05 | 0 |
| 12 | 1.95E-05 | 0.012012012 |
| 3250 | 1.94E-05 | 0.021021021 |
| 4481 | 1.90E-05 | 0.003012048 |
| 23435 | 1.90E-05 | 0.03003003 |
| 274 | 1.90E-05 | 0.015015015 |
| 3949 | 1.88E-05 | 0.006006006 |
| 1601 | 1.88E-05 | 0.015015015 |
| 1272 | 1.87E-05 | 0.045045045 |
| 7345 | 1.87E-05 | 0.018018018 |
| 325 | 1.86E-05 | 0.033033033 |
| 836 | 1.86E-05 | 0.024024024 |
| 259 | 1.85E-05 | 0.012012012 |
| 3990 | 1.84E-05 | 0.015015015 |
| 2192 | 1.84E-05 | 0.03003003 |
| 308 | 1.83E-05 | 0.006006006 |
| 718 | 1.83E-05 | 0.009009009 |
| 4217 | 1.82E-05 | 0.009009009 |
| 5176 | 1.82E-05 | 0.006006006 |
| 7436 | 1.81E-05 | 0.039039039 |
| 56975 | 1.81E-05 | 0.03003003 |
| 2 | 1.80E-05 | 0.042042042 |
| 7374 | 1.80E-05 | 0.009009009 |
| 64837 | 1.79E-05 | 0.024024024 |
| 1325 | 1.78E-05 | 0.009009009 |
| 10128 | 1.78E-05 | 0.006006006 |
| 5664 | 1.78E-05 | 0.021021021 |
| 291 | 1.77E-05 | 0.039039039 |
| 9423 | 1.77E-05 | 0.024024024 |
| 10717 | 1.76E-05 | 0.03003003 |
| 338 | 1.76E-05 | 0.039039039 |
| 439 | 1.76E-05 | 0.003003003 |
| 3033 | 1.75E-05 | 0 |
| 1558 | 1.74E-05 | 0.015015015 |
| 54536 | 1.70E-05 | 0.009036145 |
| 811 | 1.70E-05 | 0.027027027 |
| 2023 | 1.69E-05 | 0.027027027 |
| 7042 | 1.65E-05 | 0.036036036 |
| 7040 | 1.58E-05 | 0.03003003 |
| 319 | 1.56E-05 | 0.021021021 |
| 53358 | 1.56E-05 | 0.018018018 |
| 2934 | 1.54E-05 | 0.012012012 |
| 54331 | 1.54E-05 | 0.027027027 |
| 1996 | 1.54E-05 | 0.024024024 |
| 1504 | 1.53E-05 | 0.012048193 |
| 84894 | 1.53E-05 | 0.015015015 |
| 183 | 1.52E-05 | 0.03003003 |
| 11154 | 1.52E-05 | 0.015015015 |
| 89953 | 1.51E-05 | 0.033033033 |
| 3674 | 1.50E-05 | 0.012012012 |
| 84570 | 1.50E-05 | 0.027027027 |
| 4853 | 1.49E-05 | 0.018018018 |
| 6622 | 1.49E-05 | 0.015015015 |
