## Supplementary Table 3 for "Identifying Alzheimer’s disease-associated genes using PhenoGeneRanker"

**Supplementary Table 3:** Full list of GO Biological Process enrichment of top-ranked 200 significant genes.

| **Gene Set** | **Description** | **Size** | **Expect** | **Ratio** | **P Value** | **FDR** |
| --- | --- | --- | --- | --- | --- | --- |
| GO:0034368 | protein-lipid complex remodeling | 34 | 0.41208 | 41.254 | 2.50E-24 | 1.13E-20 |
| GO:0034369 | plasma lipoprotein particle remodeling | 34 | 0.41208 | 41.254 | 2.50E-24 | 1.13E-20 |
| GO:0034367 | protein-containing complex remodeling | 37 | 0.44844 | 37.909 | 1.65E-23 | 4.96E-20 |
| GO:0042982 | amyloid precursor protein metabolic process | 72 | 0.87265 | 22.919 | 3.10E-22 | 6.98E-19 |
| GO:0071825 | protein-lipid complex organization | 53 | 0.64236 | 28.022 | 6.35E-22 | 1.14E-18 |
| GO:0097006 | regulation of plasma lipoprotein particle levels | 76 | 0.92113 | 21.713 | 1.04E-21 | 1.56E-18 |
| GO:0071827 | plasma lipoprotein particle organization | 50 | 0.606 | 28.053 | 8.93E-21 | 1.15E-17 |
| GO:0050435 | amyloid-beta metabolic process | 51 | 0.61812 | 27.503 | 1.33E-20 | 1.49E-17 |
| GO:0034205 | amyloid-beta formation | 43 | 0.52116 | 30.701 | 2.30E-20 | 2.30E-17 |
| GO:0006869 | lipid transport | 417 | 5.0541 | 6.9251 | 8.65E-20 | 7.78E-17 |
| GO:0010876 | lipid localization | 465 | 5.6358 | 6.3877 | 3.49E-19 | 2.85E-16 |
| GO:0006508 | proteolysis | 1498 | 18.156 | 3.4149 | 8.69E-19 | 6.52E-16 |
| GO:0042987 | amyloid precursor protein catabolic process | 54 | 0.65448 | 24.447 | 1.63E-18 | 1.13E-15 |
| GO:0034375 | high-density lipoprotein particle remodeling | 16 | 0.19392 | 56.724 | 2.58E-18 | 1.66E-15 |
| GO:0034370 | triglyceride-rich lipoprotein particle remodeling | 12 | 0.14544 | 68.756 | 3.50E-18 | 1.97E-15 |
| GO:0034372 | very-low-density lipoprotein particle remodeling | 12 | 0.14544 | 68.756 | 3.50E-18 | 1.97E-15 |
| GO:0043691 | reverse cholesterol transport | 17 | 0.20604 | 53.387 | 7.23E-18 | 3.83E-15 |
| GO:0030301 | cholesterol transport | 111 | 1.3453 | 14.123 | 6.95E-17 | 3.47E-14 |
| GO:0015918 | sterol transport | 124 | 1.5029 | 12.642 | 5.96E-16 | 2.82E-13 |
| GO:0033344 | cholesterol efflux | 57 | 0.69084 | 20.265 | 4.48E-15 | 2.01E-12 |
| GO:0006641 | triglyceride metabolic process | 104 | 1.2605 | 13.487 | 7.06E-15 | 3.03E-12 |
| GO:0006897 | endocytosis | 687 | 8.3265 | 4.4437 | 1.39E-14 | 5.70E-12 |
| GO:0006638 | neutral lipid metabolic process | 131 | 1.5877 | 11.337 | 2.57E-14 | 1.01E-11 |
| GO:0034381 | plasma lipoprotein particle clearance | 40 | 0.4848 | 24.752 | 2.96E-14 | 1.11E-11 |
| GO:0051604 | protein maturation | 525 | 6.363 | 5.029 | 3.70E-14 | 1.33E-11 |
| GO:0006898 | receptor-mediated endocytosis | 253 | 3.0664 | 7.5007 | 5.16E-14 | 1.79E-11 |
| GO:1905952 | regulation of lipid localization | 162 | 1.9635 | 9.6768 | 9.16E-14 | 3.05E-11 |
| GO:0016485 | protein processing | 238 | 2.8846 | 7.6268 | 1.34E-13 | 4.32E-11 |
| GO:1905918 | regulation of CoA-transferase activity | 8 | 0.096961 | 72.194 | 2.73E-13 | 8.19E-11 |
| GO:1905920 | positive regulation of CoA-transferase activity | 8 | 0.096961 | 72.194 | 2.73E-13 | 8.19E-11 |
| GO:0006639 | acylglycerol metabolic process | 130 | 1.5756 | 10.789 | 3.18E-13 | 9.24E-11 |
| GO:0015850 | organic hydroxy compound transport | 283 | 3.43 | 6.7056 | 5.56E-13 | 1.54E-10 |
| GO:1990000 | amyloid fibril formation | 38 | 0.46056 | 23.884 | 5.63E-13 | 1.54E-10 |
| GO:0065005 | protein-lipid complex assembly | 29 | 0.35148 | 28.451 | 8.88E-13 | 2.35E-10 |
| GO:0033700 | phospholipid efflux | 14 | 0.16968 | 47.147 | 1.14E-12 | 2.92E-10 |
| GO:0051049 | regulation of transport | 1542 | 18.689 | 2.7824 | 4.55E-12 | 1.14E-09 |
| GO:0042981 | regulation of apoptotic process | 1392 | 16.871 | 2.9044 | 4.86E-12 | 1.18E-09 |
| GO:0008203 | cholesterol metabolic process | 134 | 1.6241 | 9.8517 | 6.76E-12 | 1.60E-09 |
| GO:1902003 | regulation of amyloid-beta formation | 35 | 0.4242 | 23.574 | 7.64E-12 | 1.75E-09 |
| GO:0032879 | regulation of localization | 1940 | 23.513 | 2.5093 | 7.78E-12 | 1.75E-09 |
| GO:0006915 | apoptotic process | 1845 | 22.362 | 2.549 | 1.08E-11 | 2.32E-09 |
| GO:0051241 | negative regulation of multicellular organismal process | 1085 | 13.15 | 3.1938 | 1.19E-11 | 2.38E-09 |
| GO:0022008 | neurogenesis | 1666 | 20.192 | 2.4762 | 8.41E-10 | 1.02E-07 |
| GO:0045184 | establishment of protein localization | 1648 | 19.974 | 2.4031 | 5.43E-09 | 5.15E-07 |
| GO:0006629 | lipid metabolic process | 1325 | 16.059 | 2.6153 | 5.57E-09 | 5.22E-07 |
